# AIO Part I: Interpretable ADMET Prediction for Cyclic Peptides and Macrocycles

**DOI:** 10.64898/2026.11.18.767182

**Authors:** Sicheng Wen, Shu Wang, Yue Qian

**Affiliations:** MARS Department, Viva Biotech (Shanghai) Limited, 735 Ziping Road, Pudong New District, Shanghai, 201318, China

**Keywords:** Macrocycles, Cyclic Peptides, ADMET, Multimodal Learning, Interpretability

## Abstract

Macrocycles and cyclic peptides are increasingly important in beyond-rule-of-five (bRo5) drug discovery, yet their absorption, distribution, metabolism, excretion, and toxicity (ADMET) properties remain difficult to predict owing to sparse data, complex topology, and environment-dependent conformations. Here, we present AIO (All-in-One), an interpretable multimodal ADMET framework combining topology-preserving molecular segmentation with solvent-aware contrastive pretraining. Across repeated cross-validation, AIO performed strongly across permeability, solubility, unbound fraction, clearance, half-life, bioavailability, and toxicity endpoints, achieving Pearson R = 0.87 for PAMPA permeability and AUROC = 0.91 for bioavailability. Case studies of recently approved macrocyclic and cyclic-peptide drugs showed agreement between predictions and reported pharmacokinetic profiles, while segment-level attribution highlighted molecular regions associated with predicted ADMET bottlenecks. Despite pretraining exclusively on macrocycles, AIO also transferred competitively to the small-molecule PharmaBench dataset. These results establish a promising foundation for interpretable ADMET prediction across diverse molecular modalities in drug discovery.

## 1. Introduction

Drug discovery is increasingly expanding beyond conventional rule-of-five (Ro5) chemical space into beyond-rule-of-five (bRo5) space, encompassing structurally complex modalities capable of engaging targets that are difficult to address with traditional small molecules ^1,2^. Macrocycles and cyclic peptides are particularly attractive because their larger interaction surfaces and conformationally constrained architectures can enable modulation of protein-protein interactions, shallow binding pockets, and other challenging targets ^1,3^. Recent clinical progress further demonstrates that such molecules can achieve drug-like systemic exposure and, in some cases, oral administration. In 2026, the FDA approved three notable orally bioavailable macrocyclic drugs, including icotrokinra, a peptide antagonist of the IL-23 receptor ^4–6^; enlicitide (MK-0616), the first oral PCSK9 inhibitor ^7,8^; and daraxonrasib, a beyond-rule-of-five (bRo5) macrocycle for metastatic pancreatic cancer ^9–11^. These advances highlight the growing therapeutic relevance of bRo5 chemical space and the need for predictive ADMET approaches tailored to these modalities.

Current computational ADMET models have largely been developed for such requirements within conventional small-molecule chemical space ^12,13^. However, commonly used datasets are relatively small or originate from historical collections ^14–16^, limiting their representation of contemporary drug-like chemical diversity. More recent resources such as PharmaBench ^17^ have substantially improved open ADMET benchmarking, assembling more than 52,482 measurements across 11 endpoints, but remain centered primarily on conventional small molecules instead of bRo5 modalities. Macrocycles are therefore underrepresented in existing ADMET datasets and benchmarks.

Beyond limited data availability, model development is challenged by the incomplete mechanistic understanding of macrocycle pharmacokinetics. Their high molecular weight, large polar surface area, and conformational complexity place them outside the applicability domain of conventional small-molecule models. Although molecular chameleonicity can help some bRo5 macrocycles reconcile membrane permeability and target recognition ^18,19^, its structural determinants remain difficult to quantify and predict. Consequently, models developed for conventional small molecules may not adequately represent the environment-dependent behavior governing macrocycle exposure.

To capture these environment-dependent behaviors computationally requires molecular representations that adequately describe the structural features unique to macrocycles. However, atom-level graphs provide a general description of chemical structure but do not explicitly encode the modular organization of macrocycles ^20,21^, whereas peptide sequence representations are poorly suited to noncanonical residues, nonpeptidic macrocycles, branched structures, multiple cyclic systems, and complex linkages ^22,23^. Fragment-based representations offer an intermediate level of abstraction, but existing approaches were largely developed for conventional small molecules ^24,25^ and do not explicitly preserve higher-order macrocyclic topology. ^26^ To our knowledge, no unified segmentation framework provides a chemically interpretable decomposition across conventional small molecules and diverse macrocyclic architectures, including branched, bridged, and multicyclic systems.

Here, we develop AIO, a multimodal framework for macrocycle ADMET prediction integrating macrocyclic segmentation with chameleonicity-guided representation learning. Macrocyclic structures are decomposed into reusable chemical segments while explicitly preserving cyclic paths, branches, linkages, bridged systems, and multicyclic topologies. To capture molecular chameleonicity, we introduce contrastive pretraining across conformational ensembles in polar and nonpolar environments, together with physicochemical objectives describing three-dimensional polarity and related structural properties. The learned representation is subsequently transferred across a broad panel of ADMET endpoints, while the segment-level formulation enables perturbation-based attribution to prioritize structural regions for medicinal-chemistry optimization. We then evaluate AIO through internal validation and external case studies, segment-level interpretation, and transfer to conventional small-molecule ADMET prediction. Despite macrocycle-only pretraining and vocabulary mismatch, the learned representation remained competitive on PharmaBench. Collectively, these results support AIO as a unified framework linking macrocycle-aware representation, environment-responsive conformational learning, broad ADMET prediction, and interpretable optimization in bRo5 chemical space.

## 2. Methods

### 2.1 Definition of macrocycles

For the purposes of this study, macrocycles were defined as molecules containing at least one ring of 12 or more heavy atoms ^27^, as determined using RDKit ring perception. When multiple qualifying rings were identified, the largest was selected for subsequent analysis. The study was restricted to compounds with molecular weights between 350 and 1,800 Da. Porphyrinoid-like structures were excluded if the selected macrocycle was associated with at least four overlapping fused-ring systems and at least 80% of these systems contained nitrogen, oxygen, or sulfur. Cyclic peptides, including bridged peptides, and other amide-containing macrocycles were retained given they met the same ring-size and molecular-weight criteria. Before analysis, salts were removed and invalid or duplicate molecular records were excluded.

### 2.2 Curation of macrocyclic ADMET data

To construct the dataset for macrocycle ADMET prediction, we leveraged macrocyclic structures from public databases for pretraining and compiled experimentally measured ADMET data from literature and patents for downstream fine-tuning.

For pretraining, we assembled a structurally diverse collection of cyclic peptides and macrocycles from four recently curated public resources. CyclicPepedia ^28^ and Macrocycle-DB ^29^ originally contained 8,744 cyclic peptides and 45,525 macrocycles, respectively, whereas NPMMP-DB ^30^ and CycPeptMPDB ^31^ contributed 4,216 non-peptidic macrocycles and 7,334 cyclic peptides, respectively. These numbers represent the original database sizes before processing with our macrocycle curation pipeline. Following structure standardization, filtering, deduplication, and application of the macrocycle selection criteria, 54,842 molecular records were retained for pretraining. Although NPMMP-DB and CycPeptMPDB contain permeability-related measurements, their endpoint annotations were excluded from the pretraining objective, and only molecular structures were retained to prevent leakage of permeability labels into downstream supervised tasks.

For fine-tuning, we curated experimentally labelled macrocyclic ADMET datasets from the literature and patents. The collected endpoints spanned in vitro measurements, including solubility, permeability, and hepatic stability, as well as in vivo pharmacokinetic measurements, including terminal half-life, total clearance, and bioavailability. Data curation followed a common workflow across endpoints. Records were first extracted with their molecular structures, experimental values, and available metadata. Structures were standardized and filtered according to the macrocycle selection criteria, after which measurements were assigned to defined ADMET endpoints and annotated according to assay conditions, biological matrix, species, and administration route where available. Experimental values were converted to common units and transformed where appropriate. Duplicate, conflicting, censored, and incomplete records were subsequently handled according to endpoint-specific rules. After quality control, 20,416 macrocyclic ADMET data points were retained for downstream fine-tuning. **Table 1** summarizes the number of data points available for each endpoint, stratified by species. Detailed processing rules and endpoint-specific caveats are provided in the Supplementary Information.

**Table 1.** Overview of the macrocyclic ADMET datasets. Data for monkeys, dogs, and cattle are not shown because of the small sample sizes but will be highlighted if used in subsequent analyses.

| Property/Assay | Size | Human | Rat | Mouse | Units | Task |
|---|---|---|---|---|---|---|
| Solubility | 423 | - | - | - | μg/mL | Reg. |
| PAMPA | 10,108 | - | - | - | cm/s | Reg. |
| Caco-2 (A to B) | 1,767 | - | - | - | cm/s | Reg. |
| Caco-2 (B to A) | 389 | - | - | - | cm/s | Reg. |
| Caco-2 (efflux ratio) | 423 | - | - | - | N/A | Reg. |
| RRCK | 165 | - | - | - | cm/s | Reg. |
| MDCK | 63 | - | - | - | cm/s | Reg. |
| Plasma unbound fraction (f<sub>u</sub>) | 522 | 268 | 107 | 127 | % | Cls. |
| Plasma/serum half-life<sup>a</sup> | 736 | 297 | 192 | 220 | h | Cls. |
| Liver microsomal clearance (CL_int) | 1088 | 789 | 91 | 168 | μL/min/mg | Reg. |
| Vd at steady state<sup>b</sup> | 650 | 47 | 293 | 216 | L/kg | Cls. |
| Terminal half-life<sup>c</sup> | 756 | 125 | 371 | 130 | h | Cls. |
| Total clearance (CL_tot)<sup>c</sup> | 1122 | 41 | 604 | 336 | mL/min/kg | Cls. |
| Bioavailability (F) | 605 | 44 | 329 | 119 | % | Cls. |
| HC50 | 361 | - | - | - | μmol/L | Cls. |
| CC50<sup>d</sup> | 1238 | - | - | - | μmol/L | Cls. |

<sup>a</sup> In-vitro experiment to evaluate the stability of molecules in plasma; <sup>b</sup> Volume of distribution at steady state; <sup>c</sup> Terminal Vd at steady state, terminal half-life and total clearance are in-vivo measurements. Apparent clearance (CL/F) values were excluded because CL cannot be determined without an intravenous reference; <sup>d</sup> CC50 is further classified as CC50 for kidney-derived cells, hepatic cells and lymphoid and hematopoietic cells

### 2.3 Segmentation

Molecular fragmentation and substructure-based tokenization facilitate representation learning for molecular generation and ADMET prediction. Existing approaches, including GraphFP ^25^, HFST ^32^, FARM ^33^, and SimSGT ^24^, have primarily focused on conventional small-molecule representation learning, with limited explicit consideration of macrocycle segmentation. To our knowledge, segmentation frameworks explicitly tailored to diverse macrocyclic topologies remain limited. We therefore developed a macrocycle-focused, two-round segmentation framework that also accommodates small molecules by combining chemistry-informed base segmentation with data-driven vocabulary refinement. The first round opened segmentable macrocycles, defined chemically meaningful fragment boundaries, and consolidated small fragments, whereas the second refined the resulting vocabulary using fragment frequencies.

During the first-round base segmentation, macrocycles were opened at bonds consistent with plausible ring-forming reactions. Candidate disconnections included amide and disulfide bonds, with non-aromatic C–C, C–N, C–O, C–S, and C=N bonds considered as fallbacks. A bond was cleaved only if it belonged to a macrocycle and its removal reduced the number or size of macrocyclic rings. Ring information was then recomputed, and all remaining rings were protected during subsequent segmentation. Specific rules for functional groups were applied to isolate sulfonamides, ureas/thioureas, esters, conjugated dienes, hydroxy-imines/oximes, disulfides, amides, and linkers. Small terminal fragments were subsequently merged with adjacent segments without crossing protected functional-group or ring boundaries.

During the second-round vocabulary refinement, segments occurring fewer than 10 times were considered for further decomposition. Candidate splits were generated by cleaving one or two non-aromatic, non-ring single bonds and were retained when every resulting fragment contained at least three heavy atoms. Splits producing frequently observed child fragments were favored, whereas additional cuts and excessive fragmentation were penalized. Split rules were learned iteratively until no new rules were identified and were then applied recursively. Atom mappings were recovered for refined fragments whenever possible, after which the segment graph and its topological annotations were reconstructed using the same virtual-closure procedure.

To annotate macrocyclic topology, cleaved opening bonds were reintroduced as virtual closure edges in the segment graph, and the corresponding cyclic paths were reconstructed. A single cyclic system was annotated as simple or branched, whereas two or more distinct cyclic systems were annotated as multicyclic, with their connecting paths and branches recorded explicitly. Bridged topologies were identified separately when overlapping cycles or additional paths connected two positions within the same cyclic system. Structures containing both bridged and independent cyclic systems were classified as mixed topologies. The resulting topology annotations were serialized using a custom notation inspired by Hierarchical Editing Language for Macromolecules (HELM) ^34^, with explicit descriptions of cyclic organization, linkages, and branches. Examples of these annotations and segmentations are provided in **Figure S2**.

To prevent leakage from downstream ADMET datasets, vocabulary construction and refinement were performed exclusively on 54,842 macrocycles from the pretraining dataset. Base segmentation produced a vocabulary of 12,077 entries, including one unknown (UNK) token. Of these, 2,035 fragment types occurred at least 10 times, whereas 5,519 were singletons. The refinement procedure learned and applied 5,516 splitting rules, reducing the final vocabulary to 7,704 entries.

### 2.4 Model architecture

The framework integrates a topology-aware sequence encoder and a solvent-conditioned conformer encoder to capture molecular connectivity and solvent-dependent conformational behavior (**Figure 1 a–b**). Their representations are jointly pretrained and combined with molecular descriptors for task-specific ADMET prediction (**Figure 1 c–d**).

**Figure 1.**
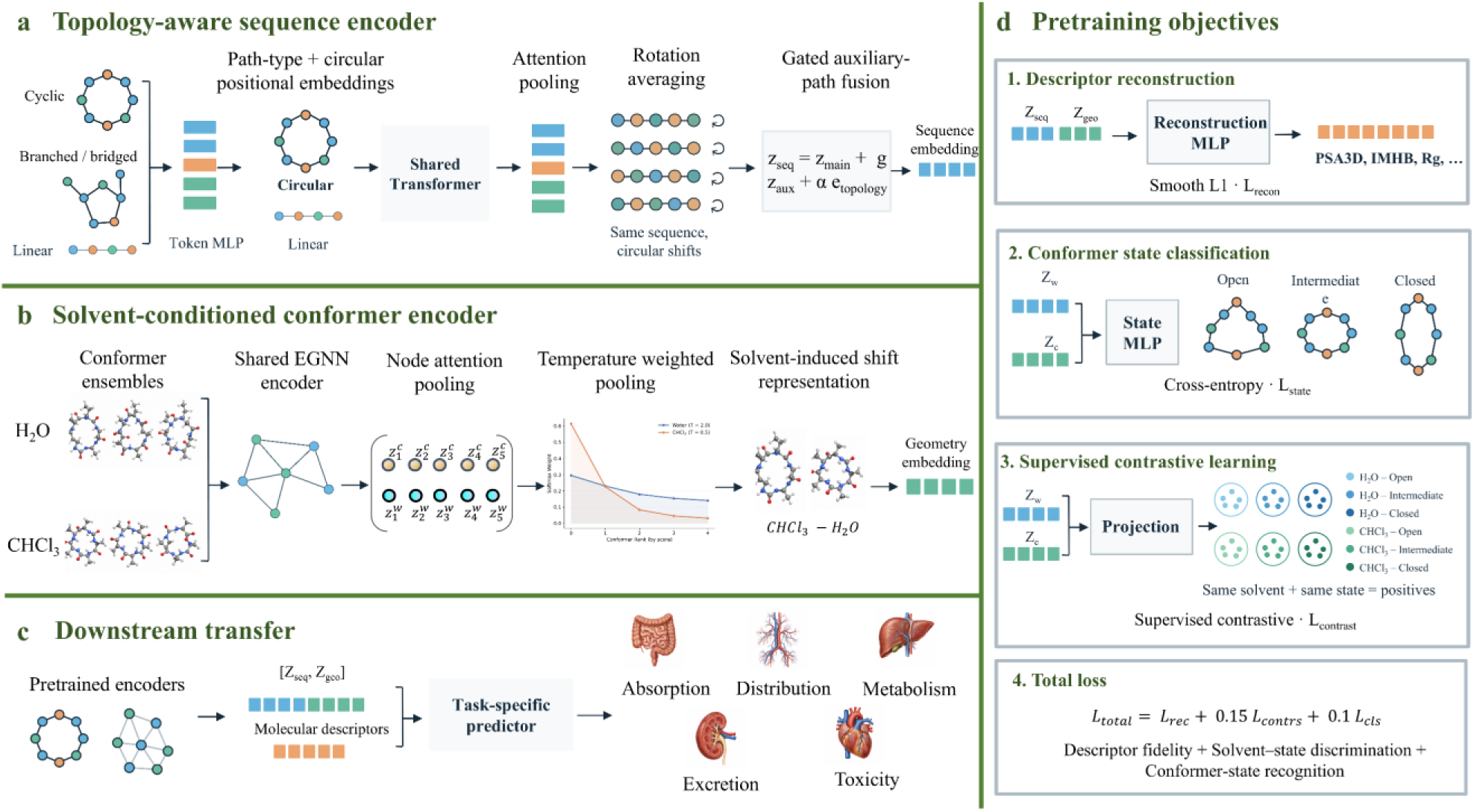
Overview of the AIO framework for ADMET prediction. (a) The topology-aware sequence encoder processes cyclic, branched or bridged, and linear paths using a shared Transformer. Circular positional embeddings encode ring periodicity, rotation averaging reduces sensitivity to the starting monomer, and gated fusion incorporates auxiliary paths. Matching colors indicate identical monomers across cyclic shifts. (b) The solvent-conditioned conformer encoder combines graph encoding, attention pooling, and temperature-weighted aggregation of water and chloroform ensembles to represent solvent-dependent conformational behavior. (c) Pretrained sequence and geometry embeddings are combined with molecular descriptors for task-specific ADMET prediction. (d) Joint pretraining integrates descriptor reconstruction, conformer-state classification, and supervised contrastive learning using solvent–state labels.

A central challenge in encoding cyclic sequences is that a cyclic path has no unique starting point. We therefore inject circular-topology information into the model through two complementary mechanisms. Circular positional embeddings encode ring periodicity, while averaging pooled representations across selected cyclic shifts reduces dependence on the arbitrary choice of sequence origin. Because these shifts preserve monomer identity, cyclic order, and connectivity, the resulting representations are encouraged to remain consistent across equivalent descriptions of the same macrocycle. Auxiliary-path information and topology embeddings additionally encode branching and bridging features. However, the current segmentation rules do not cover all macrocyclic architectures. Several macrocycles with complex fused and bridged architectures, such as systems containing three or four fused rings with an additional bridge, cannot be segmented using the current rules. These structures receive a linear-path annotation as an encoding fallback (**Figure S2**). Supporting for linear-path inputs also provides a basis for extending the framework to additional molecular modalities, including small molecules and linear peptides, although predictive performance in these modalities requires dedicated evaluation.

Furthermore, the conformer branch complements the path-based representation by capturing structural variation across ensembles in water and chloroform. Conformer sampling and ensemble construction followed the procedures described in our previous study ^35^. Aggregation within each solvent incorporates contributions from multiple conformers, while comparison between solvent-specific representations captures changes in conformational organization across the two environments. Overall, the sequence and conformer branches provide complementary descriptions of molecular topology and three-dimensional conformational behavior.

### 2.5 Training strategies and downstream fine-tuning

The model was pretrained using multitask objectives combining solvent–state supervised contrastive learning, physicochemical descriptor reconstruction, and conformational-state classification (**Figure 1 d**). Each molecule was represented by five conformers in water and five in chloroform, comprising two open, one intermediate, and two closed conformers per solvent, as described in our previous work ^35^. Combining the three conformational states with the two solvents yielded six solvent–state classes.

Supervised contrastive learning organized conformer representations according to these six classes. Within each batch, conformers sharing the same solvent–state label were treated as positive pairs, and those with different labels as negative pairs. Representations were projected into a dedicated contrastive space, normalized to unit length, and compared using cosine similarity. This objective encouraged conformers with matching solvent and state labels to occupy similar regions of the representation space. An auxiliary three-class classification task provided direct supervision for distinguishing open, intermediate, and closed conformations.

In parallel, descriptor reconstruction encouraged the learned molecular representations to retain physicochemical information. Previous studies have associated solvent-dependent changes in three-dimensional polar surface area (ΔPSA3D) with molecular permeability ^35–37^. Motivated by these findings, reconstruction targets included absolute PSA3D values in water and chloroform, thereby supervising both quantities underlying ΔPSA3D. Radius of gyration (Rg), intramolecular hydrogen bonding (IMHB), and solvent-accessible surface area (SASA) provided additional structural and physicochemical supervision. Descriptor reconstruction was optimized using the Smooth L1 loss, while conformational-state classification used cross-entropy loss.

The complete pretraining objective was defined as eqn 1

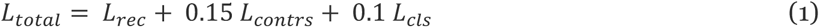

where *L_rec_* is the descriptor-reconstruction loss, *L_contrs_* is the solvent–state supervised contrastive loss, and *L_cls_* is the conformational-state classification loss. A contrastive-loss weight of 0.15 substantially improved solvent–state representation separation, with little additional gain at 0.30 (**Figure S3**). Moderate contrastive supervision also improved several solvent-specific physicochemical predictions, particularly IMHB and SASA, while largely preserving overall reconstruction performance.

Regarding data splitting, molecular connectivity was used to group compounds to minimize structural leakage. The 54,842 curated structures were partitioned into pretraining and validation sets at an 80:20 ratio. Training was performed with AdamW using a learning rate of 0.0005, weight decay of 0.01, and a batch size of 32. Training continued for a maximum of 50 epochs, with early stopping after 10 consecutive epochs without improvement in the total validation loss. Following pretraining, the pretraining-specific heads were removed. For each downstream task, the molecular encoder was initialized using the pretrained parameters. A new task-specific prediction head was introduced according to the type and structure of the endpoint labels. The output dimensionality and loss function were adapted to each task.

### 2.6 Model evaluation

Model performance was evaluated using endpoint-specific criteria designed to reflect both predictive accuracy and pharmacokinetically meaningful decision regimes. For classification endpoints, predefined thresholds were selected to represent PK regimes relevant to lead optimization and compound prioritization. Cutoffs were guided by established physiological and practical reference points, including plasma protein binding, distribution, hepatic clearance, systemic persistence, and oral exposure, while also considering the range represented in each dataset ^38,39^. Species-specific thresholds were sometimes used depending on dataset. For example, because the human total clearance dataset contains relatively few high-clearance macrocycles, thresholds were concentrated in the low-clearance range most relevant to exposure optimization. The selected thresholds and their rationales are summarized in **Table S2**.

Regression endpoints were evaluated using continuous predictions with Pearson correlation coefficient (R), mean absolute error (MAE), and root mean squared error (RMSE). Classification endpoints were evaluated at the predefined thresholds using area under the receiver operating characteristic curve (AUROC), area under the precision–recall curve (AUPRC), and balanced accuracy (BACC). The same thresholds were retained throughout model development and evaluation and were not adjusted according to test-set performance. For downstream ADMET modeling, records corresponding to the same standardized molecular structure were assigned to the same partition. For solubility, permeability, liver microsomal clearance, plasma/serum half-life and toxicity, a 20% independent fixed test set was first held out, followed by five-fold cross-validation on the remaining data for model development. For other endpoints, where reserving a fixed holdout would substantially reduce the available training data and yield an unstable estimate of performance, five-fold cross-validation with three repeats was used. Fixed test sets, when used, were excluded from model selection and hyperparameter optimization.

To further assess model behavior beyond the internal evaluation sets, five external macrocycles (icotrokinra, enlicitide, simeprevir, cyclosporine, and daraxonrasib) were used for comparison with available ADMET prediction tools. Comparisons were restricted to experimentally compatible endpoint, species, assay, and administration-route combinations. For classification endpoints, the same predefined decision regimes were used to assess agreement between predicted and experimental PK categories. Icotrokinra, enlicitide, and daraxonrasib were subsequently selected for detailed case-study analysis.

## 3. Results and discussion

### 3.1 Model performance

As shown in **Figure 2**, the model showed consistent predictive performance across ADMET endpoints, with generally good agreement between cross-validation and held-out test results. For regression tasks, permeability-related endpoints performed well. PAMPA and Caco-2 A to B reached test-set Pearson correlations of 0.87 and 0.83, respectively, while Caco-2 B to A and RRCK also showed good agreement (R = 0.74 and 0.77). Solubility and human intrinsic clearance achieved a test R of 0.75 and 0.74 on the test set respectively. AIO was further evaluated against conventional molecular-graph and Morgan-fingerprint regression baselines, as well as an equivalent model without pretraining (**Table S3)**. Similarly, classification endpoints showed strong discrimination, with AUROC values generally around 0.8–0.9. Plasma/serum half-life and HC50 achieved test AUROCs of 0.92 and 0.91, respectively, while human unbound fraction reached 0.85. The major human PK endpoints also showed strong cross-validation performance, including bioavailability (AUROC = 0.91; AUPRC = 0.96), terminal half-life (AUROC = 0.81), total clearance (AUROC = 0.88), and Vdss (AUROC = 0.85). The contribution of pretraining to classification performance was evaluated using macro-AUROC and macro-BACC across species and decision thresholds (**Table S4**). Detailed model performance for each decision threshold and species with standard deviation is reported in **Figure S4**.

**Figure 2.**
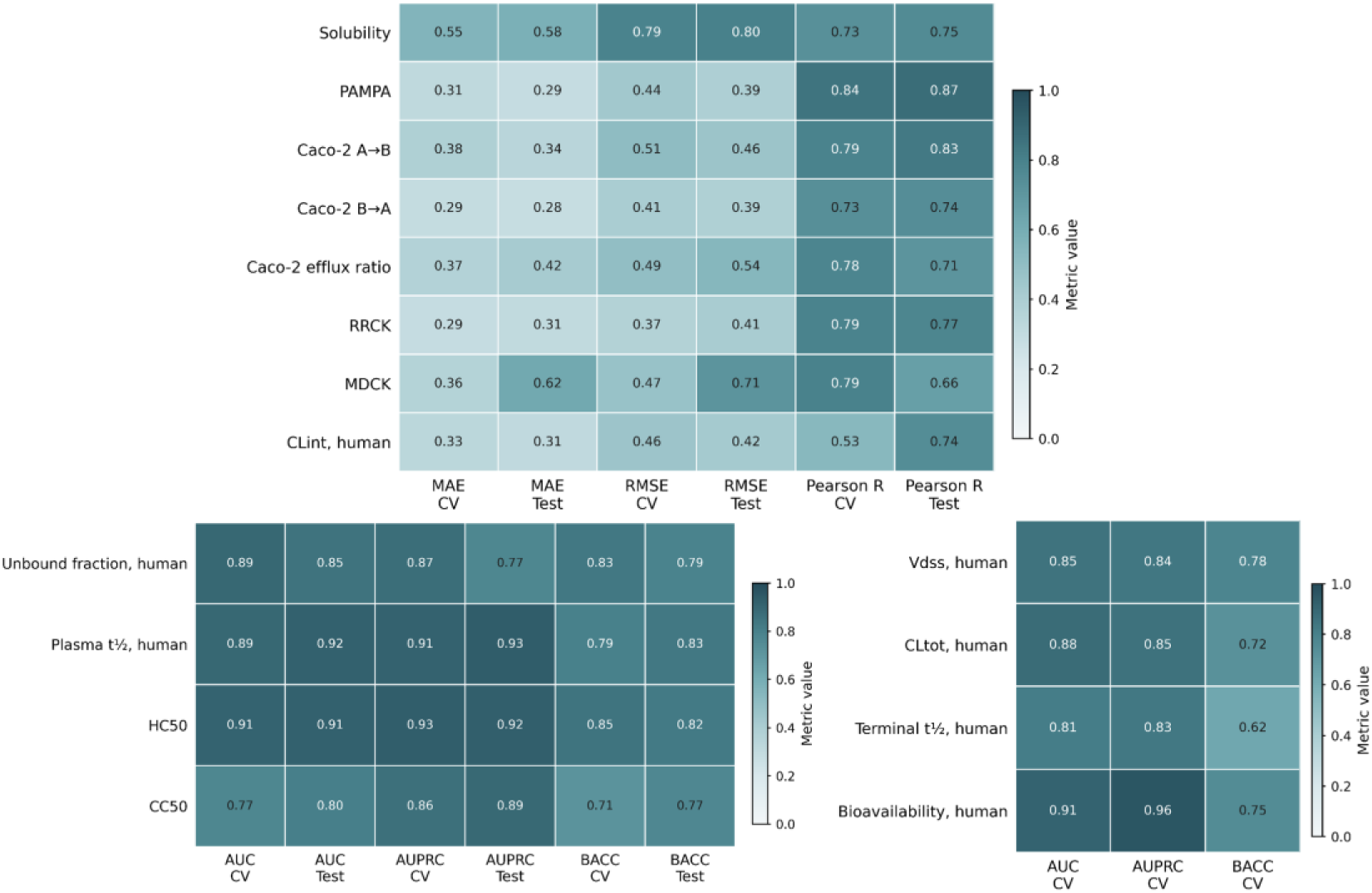
Performance of AIO across human ADMET endpoints. Regression endpoints (top) are summarized using MAE, RMSE, and Pearson correlation coefficient (R) for repeated cross-validation (CV) and held-out test sets. Classification endpoints (bottom) are evaluated using AUROC, AUPRC, and BACC. For endpoints defined using multiple decision thresholds, performance metrics were averaged across all predefined thresholds. Values represent the mean performance across human-specific tasks.

To benchmark AIO against existing ADMET prediction tools, we evaluated ADMET-AI ^40^, admetSAR 3.0^41^, ADMETlab 3.0 ^42^, ADMETPred ^43^, DeepPK ^44^, pepADMET ^45^, pkCSM ^46^, MFPK ^47^, and SwissADME ^48^ on five external compounds, including icotrokinra, enlicitide, simeprevir, cyclosporine, and daraxonrasib. Among these models, ADMET-AI, admetSAR 3.0, ADMETlab 3.0, ADMETPred, and DeepPK provided sufficient overlapping predictions across the selected endpoints for inclusion in the main comparison (**Figure 3**). Overall, AIO showed consistent agreement with the experimental profiles across permeability, plasma unbound fraction, terminal half-life, and oral bioavailability. In particular, it distinguished the very low-F profiles of icotrokinra and enlicitide from the moderate F of daraxonrasib (∼24–33%) and cyclosporine (∼33%) and the higher F of simeprevir (∼62%). As an additional bioavailability challenge case, AIO also assigned the pan-RAS inhibitor, LUNA18 ^49^, to the 20–50% F category, consistent with its experimentally observed range and further supporting its ability to distinguish favorable from poor oral exposure among cyclic peptides. Terminal half-life showed greater variability across models, reflecting its dependence on multiple PK determinants and experimental conditions. Importantly, our terminal half-life was defined following intravenous administration without an absorption phase. Following oral administration, the apparent terminal half-life may be longer when slow absorption becomes rate-limiting, as in flip-flop pharmacokinetics, which may occur in poorly permeable compounds.

**Figure 3.**
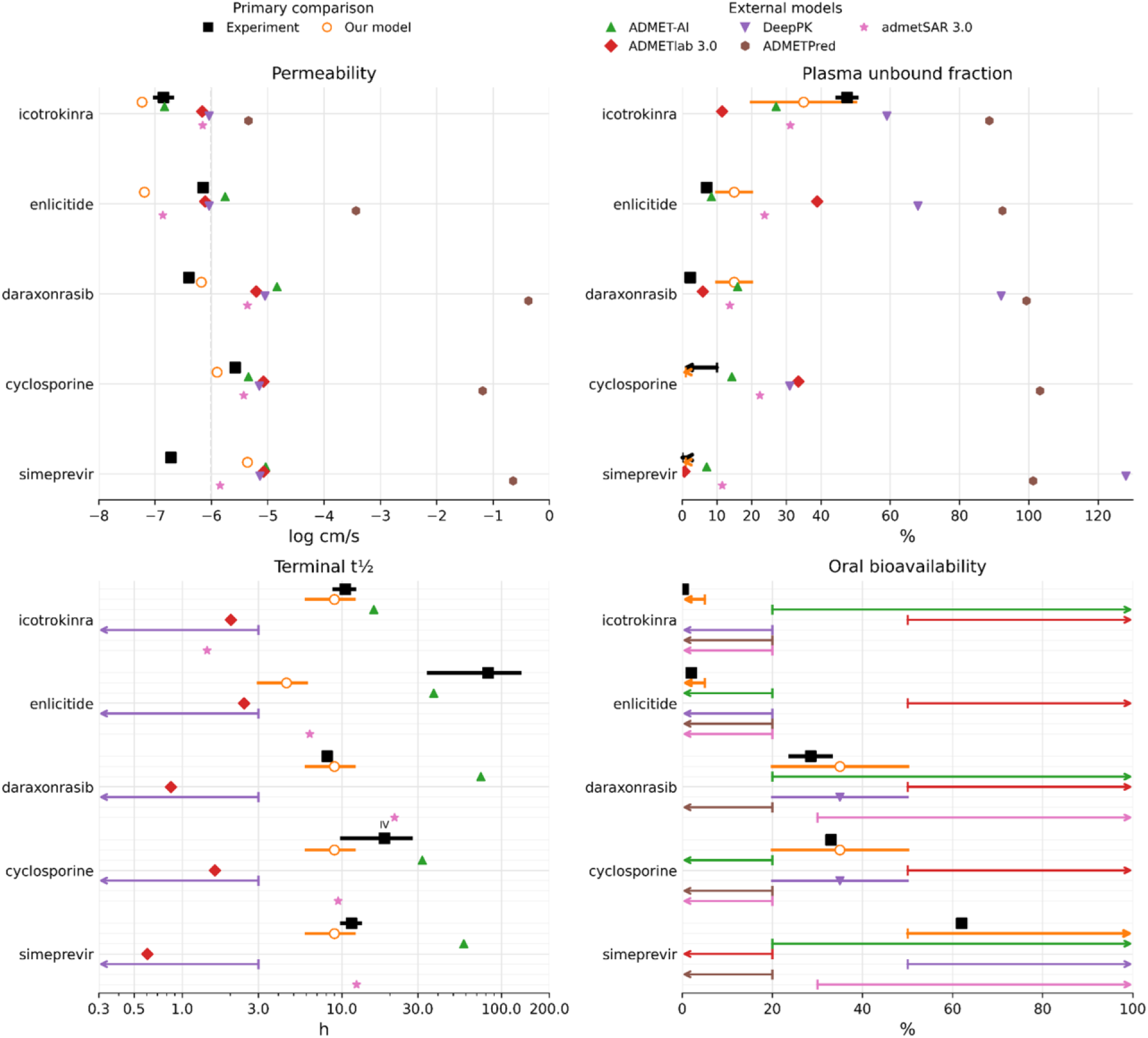
Experimental and predicted ADME profiles for five macrocyclic drugs. Experimental data represent the closest available measurements and may differ in assay system, species, route, or dose; ranges indicate reported variability across such conditions where applicable. Experimental permeability measurements include LLC-PK1 for icotrokinra, MDCK for enlicitide with cyclosporine as transporter inhibitor, and Caco-2 where available. Experimental IV terminal half-lives are indicated explicitly.

To quantify this agreement, permeability was evaluated continuously using MAE, whereas plasma unbound fraction, terminal half-life, and oral bioavailability were assessed using the predefined decision categories in **Table S2**. This category-based comparison emphasizes recovery of the experimentally supported PK state, which is directly relevant to compound prioritization and optimization; agreement was assigned when predicted and experimental values or ranges shared at least one category. In **Table 2**, AIO achieved a permeability MAE of 0.634 log cm/s, lower than admetSAR 3.0 (0.665), ADMET-AI (0.776), ADMETlab 3.0 (0.784), and DeepPK (0.826). Across the three categorical endpoints, AIO achieved the highest agreement, matching 80% of 15 evaluable comparisons, compared with 53% for ADMET-AI, 47% for admetSAR 3.0, 43% for DeepPK, 38% for ADMETPred, and 13% for ADMETlab 3.0.

**Table 2.** Comparison of model performance for permeability prediction and predefined category agreement for unbound fraction (fu), terminal half-life, and oral bioavailability (F).

| Model | Permeability MAE (n = 5)<sup>a</sup> | Valid comparisons (n/N)<sup>b</sup> | Category agreement rate |
|---|---|---|---|
| ADMETlab 3.0 | 0.784 | 15/15 | 0.13 |
| ADMETPred | 4.114 | 8/15 | 0.38 |
| DeepPK | 0.826 | 14/15 | 0.43 |
| admetSAR 3.0 | 0.665 | 15/15 | 0.47 |
| ADMET-AI | 0.776 | 15/15 | 0.53 |
| **AIO** | **0.634** | 15/15 | **0.80** |

<sup>a</sup> Permeability MAE is calculated as the mean distance between predicted and experimental points or ranges; overlapping values or ranges receive zero error. <sup>b</sup> Valid comparisons (n/N) indicate evaluable comparisons out of 15 possible molecule–property pairs. Category agreement rate is the proportion of valid comparisons sharing at least one predefined category. Missing and invalid predictions are excluded.

Thus, although individual external models occasionally matched specific measurements more closely, AIO more consistently recovered the overall experimental PK profiles across compounds and endpoints.

The remaining tools were evaluated separately because they did not provide sufficient coverage for a consistent cross-endpoint comparison. pepADMET was retained as an additional comparator because of its peptide-oriented representation, although its applicability to heavily modified or non-peptidic macrocycles is more limited. For icotrokinra, MK-0616, and cyclosporine, pepADMET predicted oral bioavailability below 20%, correctly identifying the low-F profiles of icotrokinra and MK-0616 but underestimating cyclosporine (∼33% experimentally). It also returned identical Caco-2 (−6.82 log cm/s) and RRCK (−6.20 log cm/s) predictions for icotrokinra and MK-0616 despite their distinct structures, whereas their PAMPA predictions differed (−6.59 and −5.42 log cm/s, respectively). For icotrokinra, the predicted plasma/serum half-life of 8.6 h was lower than the experimentally demonstrated stability of at least 16 h. pkCSM and MFPK provided predictions only for simeprevir and daraxonrasib, whereas SwissADME provided only bioavailability estimates for the endpoints considered here.

### 3.2 Comprehensive and integrated PK profiling

Beyond endpoint-level predictive performance, the value of a multi-endpoint ADMET framework lies in its ability to construct a coherent pharmacokinetic profile from properties that act at different stages of drug disposition. Absorption, first-pass loss, systemic disposition, and pharmacodynamic requirements jointly shape the exposure and persistence needed for therapeutic activity.

Individual ADMET properties contribute to different stages of drug disposition and are therefore most informative when interpreted collectively. At a higher level, oral bioavailability can be viewed as the product of the fraction absorbed and the fractions escaping intestinal and hepatic first-pass loss (*F* = *F_a_* × *F_g_* × *F*_ℎ_) ^50,51^. Solubility, membrane permeability, transporter effects, gastrointestinal stability, and intestinal and hepatic first-pass clearance therefore determine how efficiently an oral dose reaches the systemic circulation. Once in the circulation, systemic exposure can be interpreted through the free-drug principle, under which unbound drug is generally available for tissue distribution, clearance, and target engagement ^52,53^. Metabolic and plasma stability, protein binding, total clearance, and distribution determine the magnitude and persistence of this exposure. Pharmacodynamic properties provide an additional layer of exposure efficiency. Potency determines the concentration required to produce a therapeutic effect, while binding kinetics can influence how long target engagement persists after plasma concentrations decline ^50^. For example, a slower dissociation rate (k_off_) can prolong target residence time and sustain pharmacological activity without necessarily increasing systemic drug exposure ^54^. Thus, therapeutic persistence can arise from both pharmacokinetic mechanisms that maintain free-drug concentration and pharmacodynamic mechanisms that maintain target engagement. Overall, these factors determine the exposure required for efficacy and ultimately influence the dose and dosing interval needed in practice.

A comprehensive ADMET profile is therefore more informative than any individual endpoint, as clinically useful profiles can arise from different combinations of absorption, stability, clearance, distribution, and potency, while apparently favorable properties may sometimes reflect limitations elsewhere. For example, low membrane permeability can limit both oral absorption and access to intracellular metabolic enzymes, resulting in low bioavailability with apparently high stability in hepatocyte assays. Cross-endpoint interpretation also depends on experimental context, including species, administration route, assay conditions, and nonspecific binding; detailed considerations are provided in **Supplementary Information**.

In the present framework, endpoint-specific predictions are interpreted collectively to identify potential PK bottlenecks and trade-offs. The following external case studies apply this profile-level view to the distinct pharmacokinetic characteristics and optimization challenges of each compound.

### 3.3 Case studies: identifying compound-specific PK bottlenecks

To demonstrate the value of interpreting ADMET predictions collectively, we selected three of the five externally benchmarked compounds, including icotrokinra, enlicitide, and daraxonrasib, for detailed case-study analysis. All three received FDA approval in 2026 ^6,11,55^ and represent distinct macrocyclic or bRo5 modalities with different therapeutic mechanisms and pharmacokinetic challenges. Icotrokinra is an orally administered peptide antagonist of the IL-23 receptor for plaque psoriasis ^4,5^, whereas enlicitide is an oral macrocyclic peptide PCSK9 inhibitor that disrupts the PCSK9–LDLR protein– protein interaction ^7,8^. Daraxonrasib (RASONQUE), approved for metastatic pancreatic adenocarcinoma, represents a mechanistically distinct macrocyclic modality in which a RAS(ON) tri-complex inhibitor acts as a molecular glue by recruiting cyclophilin A to active RAS ^9^. These three examples were used to examine how cross-endpoint ADMET patterns can reveal distinct PK bottlenecks and trade-offs, with icotrokinra examined in greatest depth because of the availability of complementary PK, structural, and patent evidence.

Across the three recently approved compounds, the independently predicted ADMET endpoints broadly reproduced their distinct experimental PK profiles (**Figure 4**). Icotrokinra and enlicitide showed pronounced absorption limitations, with very low experimental permeability and predicted oral bioavailability below 5%, consistent with their low experimental bioavailability (0.12–3.31% across preclinical species for icotrokinra and ∼2% for enlicitide) ^5,7^. Complementary PAMPA, Caco-2, RRCK, MDCK, and solubility predictions further supported an absorption-limited profile. Notably, this poor absorption was accompanied by high in-vitro stability. Icotrokinra showed prolonged plasma stability and very low hepatocyte intrinsic clearance, while enlicitide was highly resistant to digestive and proteolytic degradation and exhibited very low hepatic intrinsic clearance. These trends were reflected by predicted plasma t_1/2_ > 12 h and relatively low microsomal CLint for both compounds.

**Figure 4.**
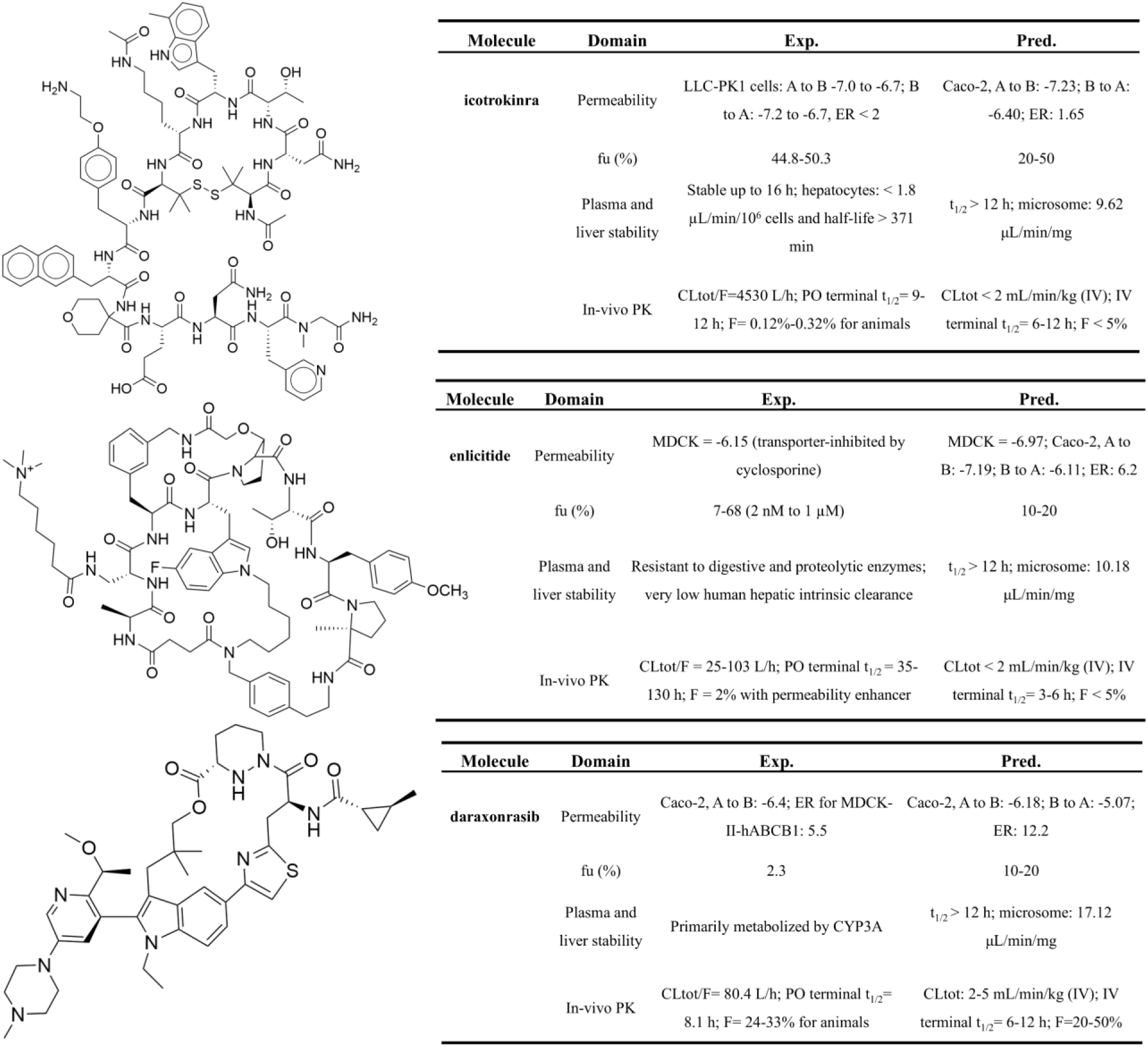
External case studies of recently approved macrocyclic drugs. Chemical structures and comparison of experimentally reported and AIO-predicted ADMET profiles for (a) icotrokinra, (b) enlicitide, and (c) daraxonrasib. The comparison includes permeability, plasma unbound fraction (fu), plasma and liver stability, and in vivo pharmacokinetic properties. Experimental values were compiled from reported preclinical studies, while predictions were generated using the corresponding AIO endpoint models.

Daraxonrasib showed a distinct profile. Although its permeability remained relatively low, the predicted Caco-2 permeability (−6.18 log cm/s) indicated a less severe barrier than for icotrokinra (−7.23) and enlicitide (−7.19), consistent with its substantially higher experimental oral bioavailability of 24–33% ^9^ and predicted range of 20–50%. Its smaller, less peptide-like structure may partly contribute to this reduced permeability constraint, although other absorption and first-pass determinants are also involved. Daraxonrasib additionally showed strong plasma protein binding (f_u_ = 2.3%), contrasting with the substantially higher unbound fraction of icotrokinra and enlicitide, while its predicted plasma stability and microsomal clearance supported sufficient metabolic stability. Collectively, these cases show that AIO captured distinct experimental PK profiles and highlight the value of interpreting permeability, bioavailability, stability, and protein binding as an integrated ADMET profile.

### 3.4 Case studies: from PK bottlenecks to structural optimization opportunities

Having identified the major PK bottlenecks at the compound level, we next asked whether segment-level interpretation could prioritize regions for subsequent medicinal-chemistry investigation and reveal potential opportunities for optimization. Icotrokinra was selected as the primary example because its low-bioavailability, absorption-limited profile provided a clear basis for linking cross-endpoint interpretation with structural attribution.

To localize structural contributions to the predicted pharmacokinetic phenotype, we applied single-segment perturbation analysis. Each predefined macrocyclic segment was individually masked and the model prediction was recalculated, with segment attribution defined as the difference between the original and masked predictions (Δ*_i_*= *y_original_* − *y_masked_*_,*i*_); for peptide-derived segments, the backbone and associated side chain were masked together whenever possible. A positive attribution therefore indicates that the segment increases the predicted output. The magnitude of Δ*_i_* reflects the sensitivity of the model prediction to that structural region. For icotrokinra, several localized regions showed substantial negative contributions to predicted oral bioavailability (**Figure 5**), with the complete segment-level attribution profile provided in **Figure S5**.

**Figure 5.**
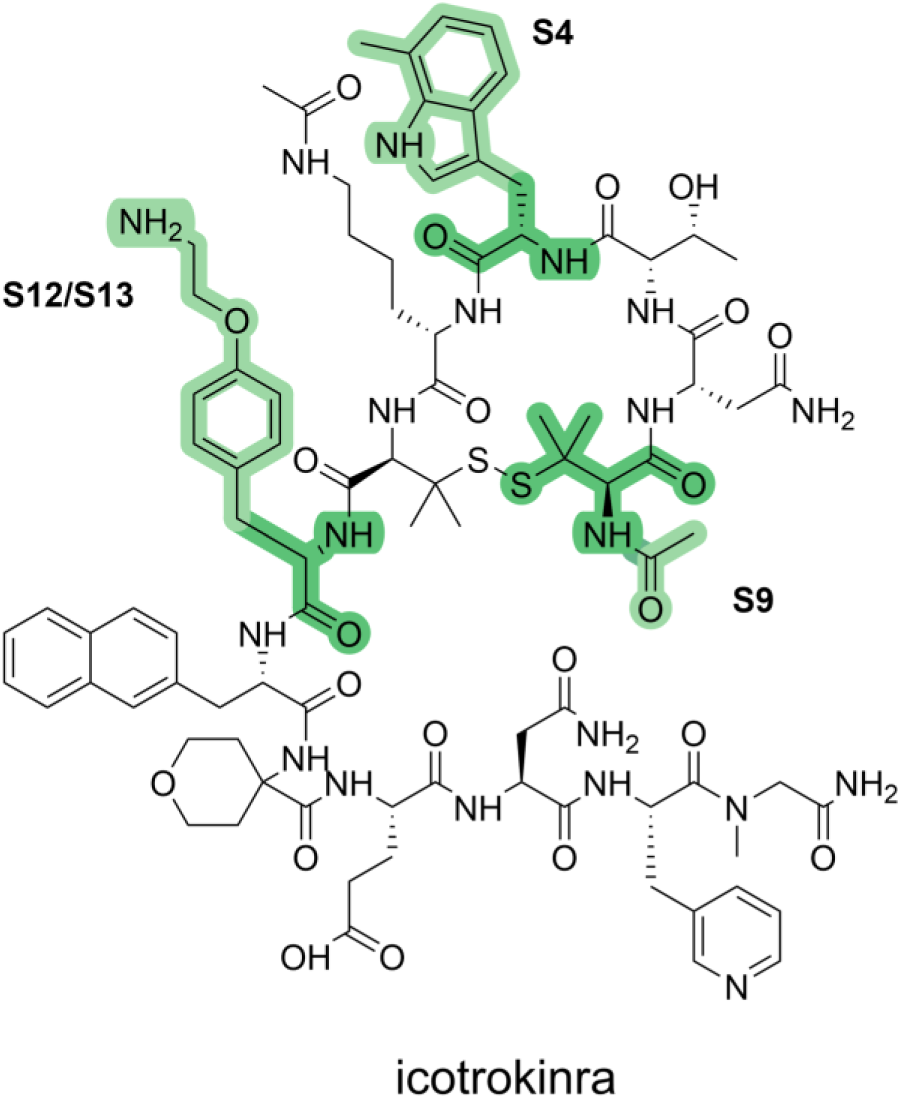
Model-highlighted structural regions of icotrokinra for further analysis. Chemical structure of icotrokinra showing segments S4, S12/S13, and S9, highlighted in green as representative regions examined in the subsequent PK and analogue analyses.

We next examined whether the regions highlighted by the model corresponded to structural positions independently explored during medicinal chemistry optimization. The disclosed analogues reflect progression from identifying activity-critical regions to optimizing sites for PK (**Figure 6**). The S4-containing region, for example, showed a substantial negative contribution to predicted oral bioavailability while modifications at this position also affected inhibitory potency. Compound 6 (WO2026057052A1) and Compound 16 (WO2024227437A1) retained activity following modification of this region, whereas compound 27 (WO2024155547A2) showed reduced potency, indicating a potential PK–activity trade-off. Similarly, the disulfide-constrained core was not considered a straightforward substitution site because changes to the cyclized scaffold may alter ring conformation and target binding ^56,57^, consistent with our molecular docking analysis (data not shown).

**Figure 6.**
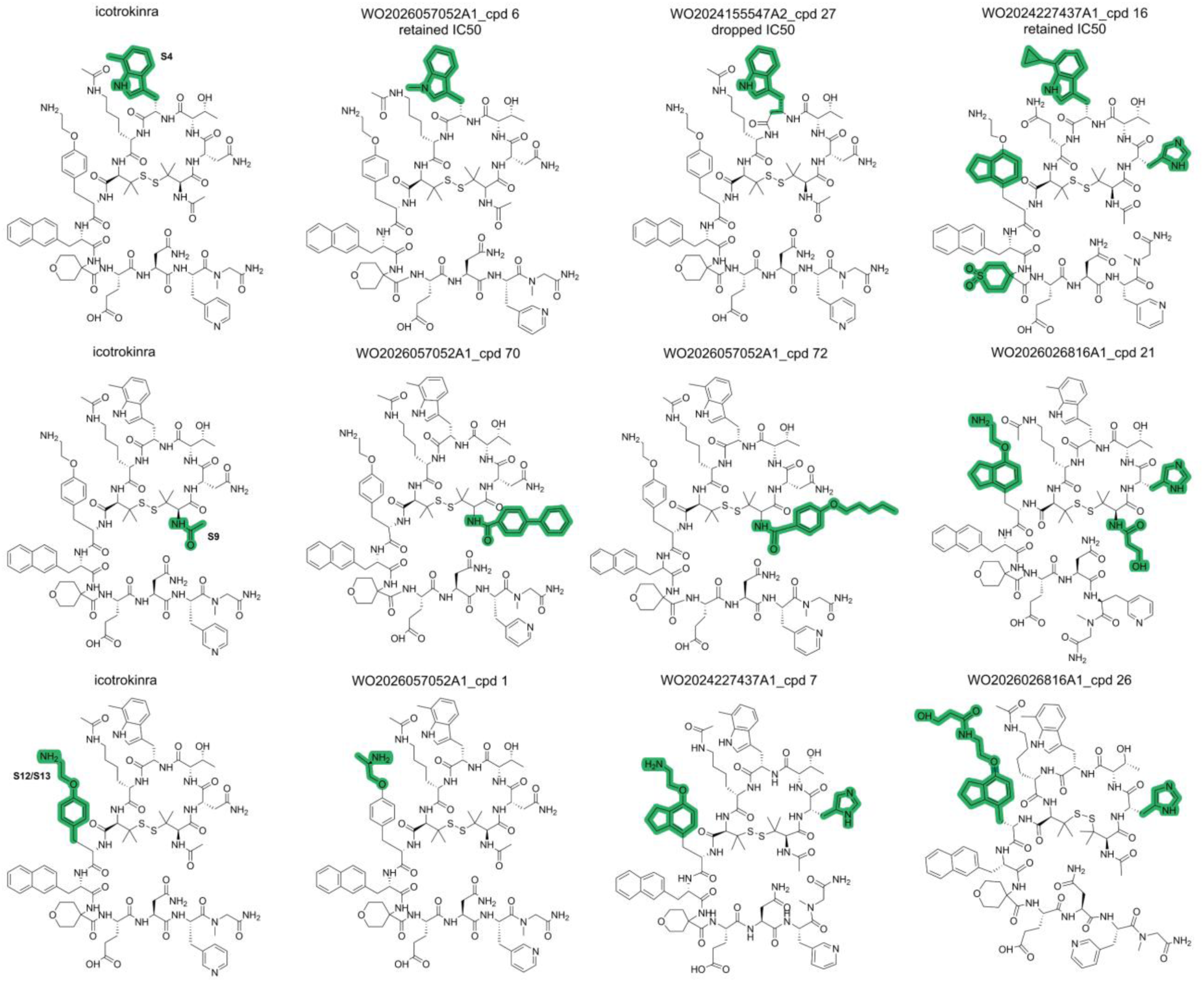
Comparison of model-prioritized icotrokinra regions with structural modifications in disclosed analogues. Icotrokinra segments highlighted by the analysis are shown alongside representative analogues reported in the indicated patents. Structural differences relative to icotrokinra are highlighted in green.

Subsequent analogues targeted other model-highlighted regions, where structural changes substantially altered PK exposure (**Table 3**). For example, compounds 1, 70, and 72 from WO2026057052A1 showed rat IV AUC_last_ values of 3,360, 3,710, and 3,530, respectively, compared with 1,910 for icotrokinra at the same IV dose of 1 mg/kg. Several analogues also showed markedly higher mouse oral exposure, including compounds 7 (WO2024227437A1), 26 and 21 (WO2026026816A1), and 16 (WO2024227437A1), with AUC_last_ values of 143–227 compared with 34.8 for icotrokinra at 10 mg/kg. Changes in these regions were also associated with altered absolute oral bioavailability; for example, compound 21 reached a reported mouse F of 0.68%, compared with 0.55% for icotrokinra, although differences in the PO/IV dose combinations limit direct quantitative comparison. Overall, these observations indicate that segment-level attribution can identify both PK-sensitive regions that are constrained by target activity and more tractable positions that have independently been exploited to modulate systemic and oral exposure.

**Table 3.**
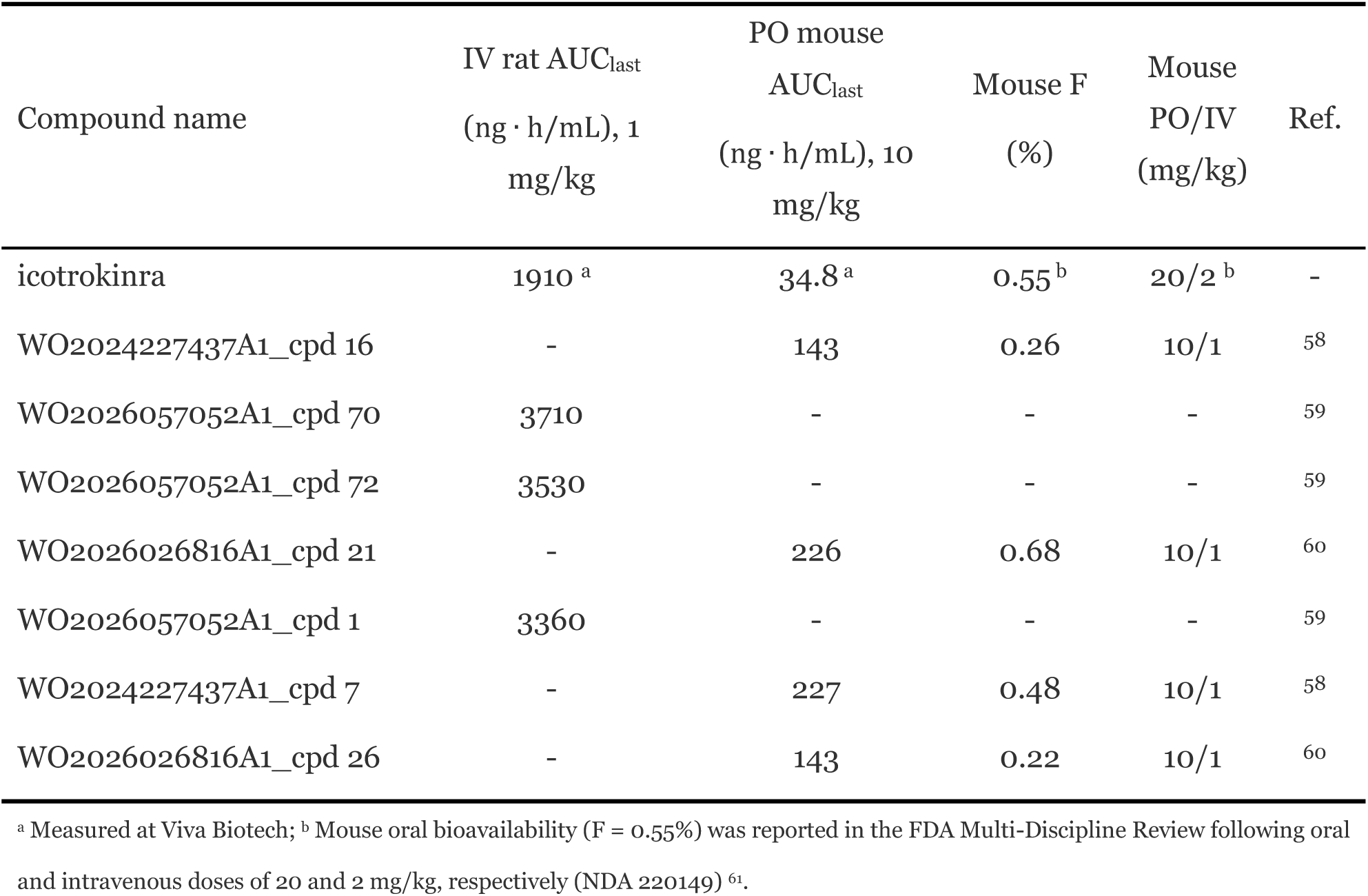
Pharmacokinetic exposure of icotrokinra and selected disclosed analogues containing modifications in model-prioritized regions. Reported rat intravenous (IV) and mouse oral (PO) AUC_last_ values are shown together with mouse oral bioavailability (F) where available. Mouse F values are accompanied by the corresponding PO/IV dose pair

### 3.5 Generalization to small-molecule ADMET prediction

To assess generalization beyond macrocyclic chemical space, we evaluated AIO using PharmaBench’s predefined scaffold split for the human liver microsomal clearance subset ^17^. The original scaffold-based train/test partition was retained, with the fixed test set kept untouched throughout model development. The PharmaBench training set was further subjected to five-fold cross-validation, where four folds were used for training and the remaining fold for validation. Despite being pretrained exclusively on macrocycles, AIO achieved performance comparable to models developed using conventional small-molecule chemical space, with Pearson R = 0.749, RMSE = 0.563, and MAE = 0.424 on the fixed test set (**Table 4**). Recent models including FragFormer ^62^ and BiScale-GTR ^63^ primarily report RMSE on PharmaBench; we additionally report Pearson R and MAE to provide complementary measures of compound-level variation and absolute prediction error.

**Table 4.** Generalization performance of AIO on the human liver microsomal clearance subset of the PharmaBench small-molecule benchmark. The pretraining domain denotes the primary chemical space used during model pretraining. Pearson R, RMSE, and MAE are reported where available; “–” indicates metrics not reported in the original study. Results for XGBoost, Uni-Mol, Transformer-M, and MPG were obtained from PharmaBench.

| Model | Pretraining domain | Pearson R | RMSE | MAE |
|---|---|---|---|---|
| XGBoost | - | 0.628 | 0.647 | 0.459 |
| Uni-mol | Small molecules | 0.722 | 0.613 | 0.426 |
| Transformer-M | Small molecules | 0.733 | 0.567 | 0.392 |
| MPG | Small molecules | 0.751 | 0.541 | 0.375 |
| FragFormer | Small molecules | - | 0.514 | - |
| BiScale-GTR | Small molecules | - | 0.501 | - |
| AIO | Macrocycles | 0.749 | 0.563 | 0.424 |

Notably, this transfer was achieved despite substantial mismatch between the macrocycle-derived segment vocabulary and conventional small-molecule chemical space. Overall, 25.5% of segment positions were mapped to the UNK token, 78.7% of molecules contained at least one unknown segment, and these assignments represented 1,002 distinct unseen segment types. The competitive performance under this setting indicates that the macrocycle-pretrained representation retains substantial transferability to conventional small molecules even when a considerable fraction of their local structural patterns is not explicitly represented in the existing vocabulary. At the same time, this incomplete vocabulary coverage suggests additional room for improvement by extending both the pretraining corpus and segment vocabulary to broader small-molecule chemical space.

## 4. Limitation and future development

Despite the overall robustness of the framework, several limitations remain. Although the ADMET datasets were curated and standardized, measurements originated from different assay formats, species, biological matrices, administration routes, and experimental conditions. Such heterogeneity is inherent to public and literature-derived pharmacokinetic data and may introduce additional variability into model training and evaluation. This is particularly relevant for macrocycles, for which available datasets remain substantially smaller and less standardized than those for conventional small molecules. The limited availability of large, consistently annotated macrocycle datasets also restricts the scale of systematic external validation. Predictions should therefore be interpreted in the context of the endpoint definitions and experimental settings represented in the training data.

Additionally, predictive performance was assessed using repeated cross-validation and held-out test sets, but out-of-distribution evaluation remained limited to a small number of external macrocycles. Segment-level interpretation was likewise evaluated retrospectively using independently disclosed patent analogues. Modifications at several model-highlighted regions were associated with improved systemic exposure and oral bioavailability, providing independent support for the relevance of the derived attributions. However, because these analogues were not prospectively designed based on model predictions, prospective experimental validation of model-guided modifications will be important for establishing the practical utility of the interpretability framework.

## 5. Conclusion

In conclusion, this study establishes a unified and interpretable ADMET framework for macrocycles and other bRo5 modalities. By combining macrocycle-aware structural representation with multi-endpoint prediction, AIO enables systematic evaluation of compounds whose pharmacokinetic behavior is difficult to capture using models developed primarily for conventional small molecules. Across internal evaluation and selected out-of-distribution macrocycles, the model showed robust performance across key properties related to absorption, systemic disposition, metabolic stability, clearance, and oral bioavailability. Integrating these endpoints provides a more complete view of PK behavior, which is particularly important for macrocycles and orally delivered peptides requiring simultaneous optimization of permeability, solubility, stability, protein binding, clearance, and exposure. Segment-level interpretation further links predicted ADMET liabilities to specific structural regions, supporting both property prediction and molecular optimization. These capabilities may help bridge increasingly complex molecular design with the pharmacokinetic requirements of successful drug development.

## Data availability

The processed pretraining data, preprocessed permeability dataset, data for the five external case studies, human liver microsomal stability data from PharmaBench, and trained models are available at Zenodo (DOI: 10.5281/zenodo.23486930). Code for figure reproduction and model inference is available at Github (https://github.com/wsicheng739/AIO-ADMET).

## Acknowledgements

We thank Dr. Xiaocong Yang for assistance with data collection and cleaning.

