## Supplementary Information for "AIO Part I: Interpretable ADMET Prediction for Cyclic Peptides and Macrocycles"

© 2026 MARS, Viva Biotech (Shanghai) Limited. All rights reserved. Visit website at www.vivabiotech.com.

### 1. Data curation and cleaning procedures

#### 1.1 Introduction

Reliable ADMET modelling requires more than cutting-edge model architectures or predictive accuracy; it also depends heavily on clearly defining what each endpoint represents, how experimental measurements are generated, and how these data should be curated and translated into modelling targets. Such practical considerations are often underreported in AI-focused studies, creating a gap between model development and the expectations of medicinal chemists and DMPK scientists. We therefore provide in this Supporting Information the endpoint definitions, data-curation procedures, experimental considerations, and rationale behind key modelling decisions for each ADMET task. Although the present study focuses on macrocycles, many of these considerations, particularly those related to endpoint definition, assay interpretation, data harmonization, and quality control, are equally relevant to conventional small-molecule ADMET modelling. By making these considerations openly available, we hope to improve the transparency and reproducibility of ADMET modelling and provide a practical reference that helps bridge the drug-discovery and AI communities.

#### 1.2 Definition of each endpoint

The endpoints included in this ADMET model can be broadly categorized into three groups, including absorption-related endpoints, in vitro stability-related endpoints, and systemic in vivo pharmacokinetic (PK) and toxicity endpoints.

Absorption-related endpoints primarily include solubility and permeability. Because of the limited availability and heterogeneity of published data, the solubility endpoint is relatively broad and contains mainly equilibrium solubility measurements obtained in aqueous buffers at approximately pH = 6.8 – 7.4. Permeability endpoints follow the assay annotations used in CycPeptMPDB [1] and include PAMPA, Caco-2, RRCK, and MDCK. PAMPA employs an artificial lipid membrane and therefore primarily measures passive membrane diffusion. RRCK, a low-efflux MDCK-derived cell line, and conventional MDCK assays provide cell-based measurements that more closely reflect transcellular permeability, although endogenous transporter activity may vary depending on the cell line. In contrast, Caco-2 cells more closely mimic the intestinal epithelial barrier and can capture both passive permeability and transporter-mediated processes. Bidirectional Caco-2 measurements are represented by apparent permeability from the apical to basolateral direction (A to B) and from the basolateral to apical direction (B to A), together with the corresponding efflux ratio.

In vitro stability-related endpoints include plasma/serum half-life and liver microsomal clearance, which probe distinct degradation and metabolic processes. Because of the limited availability of cyclic-peptide stability data, plasma and serum stability measurements were combined into a single endpoint for modeling. Both assays reflect compound stability in circulating biological fluids and are particularly relevant to cyclic peptides because proteolytic and peptidolytic cleavage can contribute to their instability. The resulting plasma/serum half-life endpoint therefore provides an indication of stability in the circulation [2]. Liver microsomal clearance, in contrast, evaluates hepatic metabolic stability and,

© 2026 MARS, Viva Biotech (Shanghai) Limited. All rights reserved. Visit website at www.vivabiotech.com.

under conventional NADPH-supplemented conditions, largely captures oxidative metabolism mediated by microsomal enzymes such as CYP450. UGT-mediated metabolism may also be evaluated when appropriate cofactors and assay conditions are included [3].

Several parameters provide important mechanistic links between in vitro measurements and systemic in vivo PK. One representative example is the plasma unbound fraction (fu), which describes the fraction of a compound that is not bound to plasma proteins and is therefore available for distribution, metabolism, elimination, and target engagement. According to the free-drug hypothesis, the unbound drug concentration is generally considered the pharmacologically and kinetically relevant fraction. Plasma protein binding is also an important component of in vitro–in vivo extrapolation (IVIVE). For example, in vivo hepatic clearance can be estimated by integrating in vitro intrinsic clearance with the unbound fraction and relevant physiological parameters using appropriate hepatic disposition models.

Systemic in vivo PK endpoints include volume of distribution at steady state (Vdss), terminal half-life, total clearance, and bioavailability. These parameters represent integrated outcomes arising from absorption, distribution, metabolism, and elimination processes. They can vary substantially across species and administration routes and, in some cases, may also exhibit dose dependence due to nonlinear PK behavior. Species and administration route were therefore incorporated as modeling conditions where relevant, whereas dose was not explicitly included because doing so would further fragment the available data and substantially increase data sparsity. Toxicity-related endpoints include HC50 and CC50. Because cytotoxic responses can depend strongly on the biological context of the assay, CC50 measurements were further categorized according to the tissue origin of the tested cell lines, including kidney-derived, hepatic, and lymphoid or hematopoietic cells.

##### 1.3 Data curation workflow

Given the diversity of experimental protocols and reporting conventions across ADMET endpoints, we use liver microsomal intrinsic clearance as a representative example to illustrate the major considerations involved in data extraction, normalization, and quality control. This endpoint requires careful distinction among related clearance measurements and experimental systems. Although assay-specific criteria differ across endpoints, the principles illustrated here, including terminology unification, unit standardization, experimental-condition filtering, and review of ambiguous records, were applied throughout the dataset.

After collecting data from patents and the literature, the raw liver-clearance records contained heterogeneous measurements, including hepatocyte clearance, liver microsomal clearance, and predicted in vivo hepatic clearance derived through in vitro–in vivo extrapolation (IVIVE). These measurements were first distinguished according to their experimental context and reporting units. Hepatocyte intrinsic clearance is commonly normalized by cell number and reported in units such as μL/min/$10^6$ ells, whereas liver microsomal intrinsic clearance is typically normalized by microsomal protein content and reported as μL/min/mg protein. In contrast, IVIVE-derived hepatic clearance is generally scaled to body weight and expressed in units such as mL/min/kg. Importantly, IVIVE-predicted hepatic clearance and experimentally measured in vivo systemic clearance may share the same

© 2026 MARS, Viva Biotech (Shanghai) Limited. All rights reserved. Visit website at www.vivabiotech.com.

units but represent fundamentally different quantities and were therefore not treated as interchangeable.

Experimental conditions were then reviewed. Liver S9 measurements were excluded because this fraction contains both microsomal and cytosolic enzymes, representing a broader metabolic system than isolated microsomes. Species annotations were standardized, retaining human, rat, mouse, monkey, dog, and bovine records where applicable. NADPH supplementation was manually verified, and only measurements obtained in its presence were retained to ensure consistency with conditions supporting CYP-mediated oxidative metabolism. Repeated measurements for the same molecule under equivalent experimental conditions were aggregated using the median. Finally, records were filtered according to the macrocycle definition adopted in this study to generate the modelling dataset.

The curation of other ADMET endpoints followed the same general principles, with endpoint-specific criteria determined by the experimental meaning of each measurement. For example, administration route was considered when interpreting in vivo clearance and terminal half-life, whereas assay system and transport direction were distinguished for permeability measurements. Nevertheless, curation cannot fully eliminate variability arising from experimental conditions or incomplete reporting. The following section therefore highlights endpoint-specific pitfalls that can complicate comparisons and biological interpretation, including those that may persist after terminology and units have been standardized.

#### 1.4 Potential pitfalls in data curation

Beyond the general curation workflow, several pitfalls can affect endpoint interpretation, training labels, and model evaluation. These issues are relevant to both AI researchers and medicinal chemists because similarly reported measurements may differ in assay design, biological system, species, administration route, or experimental conditions. **Table S1** summarizes common sources of ambiguity across ADMET endpoints, followed by detailed discussions of selected issues and their mechanistic implications. These considerations highlight limitations in measurement comparability and biological interpretation, including sources of variability that may remain after data curation.

**Table S1. Common pitfalls in ADMET data interpretation and curation and their consequences for modelling.** Selected sources of assay variability and endpoint ambiguity that can affect measurement comparability, training labels, and biological interpretation.

| Endpoint | Potential pitfall | Consequences for interpretation/modelling |
|---|---|---|
| Permeability (PAMPA, Caco-2) | Overlooking nonspecific binding, recovery, and efflux in cell-based assays | Misinterprets permeability and misguides optimization |
| Liver microsomal clearance | Concentration-dependent nonspecific binding | Alters apparent clearance and reduces label comparability |
| Plasma unbound | Concentration-dependent binding and | Increases variability in measured |

© 2026 MARS, Viva Biotech (Shanghai) Limited. All rights reserved. Visit website at www.vivabiotech.com.

| | | |
|---|---|---|
| fraction | nonspecific adsorption | unbound fraction |
| Plasma/serum stability half-life | Confusing in vitro stability with in vivo half-life | Mislabels endpoints and misleads interpretation |
| In-vivo CL and Vd | Pooling CL with CL/F or Vd with Vd/F | Confounds clearance and distribution with bioavailability |
| Terminal half-life | Pooling oral and IV measurements despite absorption-limited kinetics | Confounds absorption with elimination |
| Oral bioavailability | Overlooking food status (fasted or fed), formulation, or absolute versus relative F | Confounds experimental conditions with molecular properties |

For absorption-related endpoints, permeability measurements require particular attention to assay recovery and transport directionality. Oral drug absorption occurs predominantly in the small intestine, where membrane permeability is a determinant of systemic exposure. However, hydrophobic or highly lipophilic compounds may nonspecifically bind to membranes, cells, or assay devices during permeability experiments, resulting in poor recovery and potentially underestimated apparent permeability. Therefore, post-assay recovery should be considered whenever available [4]. In addition, cellular permeability is inherently bidirectional. Considering only A to B permeability while ignoring B to A permeability or the corresponding efflux ratio may obscure transporter-mediated effects and lead to incomplete interpretation of intestinal permeability [5]. A high ER supports directional transport, whereas identification of P-gp generally requires inhibitor or transporter-specific experiments.

Such nonspecific binding is also an important source of variability in in-vitro stability and protein-binding assays. Previous studies from AstraZeneca and Pfizer have demonstrated substantial effects of microsomal binding and protein concentration on apparent metabolic clearance [6,7]. Under the assumption that metabolism is proportional to unbound drug concentration, binding-corrected intrinsic clearance is expressed as Eqn (1)

$$\mathrm{CL_{int,u}} = \frac{CL_{\mathrm{int,app}}}{\mathrm{f_{u,mic}}} \tag{1}$$

To quantify the magnitude of variability in microsomal binding, we analyzed the dataset of fraction unbound values [8]. Among 180 compound–species groups evaluated at two or more microsomal protein concentrations, the median absolute difference in the unbound fraction ($f_{u,mic}$) between the lowest and highest tested concentrations was 0.100 (interquartile range [IQR]: 0.049–0.283; mean: 0.182). Decreasing $f_{u,mic}$ with increasing protein concentration was observed in 58.9% of groups, whereas 29.4% showed an increase and 11.7% remained approximately unchanged (absolute difference ≤0.02). The 20 groups with the largest variation exhibited absolute differences in $f_{u,mic}$ of approximately 0.45–0.81 (**Figure S1**). For example, the unbound fraction of imipramine in human microsomal incubations decreased from 0.94 at 0.025 mg/mL to 0.13 at 2 mg/mL, an approximately sevenfold difference. Chlorpromazine similarly decreased from 0.75 at 0.025 mg/mL to 0.06 at 0.71 mg/mL. These findings

© 2026 MARS, Viva Biotech (Shanghai) Limited. All rights reserved. Visit website at www.vivabiotech.com.

highlight how variation in incubation binding may complicate comparisons of apparent intrinsic clearance across assay conditions. Measuring $f_{u,mic}$ under matched incubation conditions, for example by equilibrium dialysis, allows apparent intrinsic clearance to be corrected for binding. The resulting unbound intrinsic clearance values may provide more comparable training labels for deep-learning models. However, this benefit depends on reliable $f_{u,mic}$ measurements, particularly at low values, where measurement errors can substantially affect the corrected clearance. An additional source of variability in human microsomal clearance measurements is interindividual variation in drug-metabolizing enzyme activity. Genetic polymorphisms in CYP enzymes, including CYP2D6 and CYP2C19, can substantially alter enzyme activity, and the frequencies of functional alleles vary among human populations [9]. Such donor-to-donor differences in CYP expression and activity may therefore contribute to variability in measured intrinsic clearance, particularly when single-donor or differently composed pooled human liver microsomes are used. Recording donor information, CYP phenotype or genotype, and microsomal pool composition where available may help distinguish biological variability from compound-dependent clearance and improve the consistency of training labels.

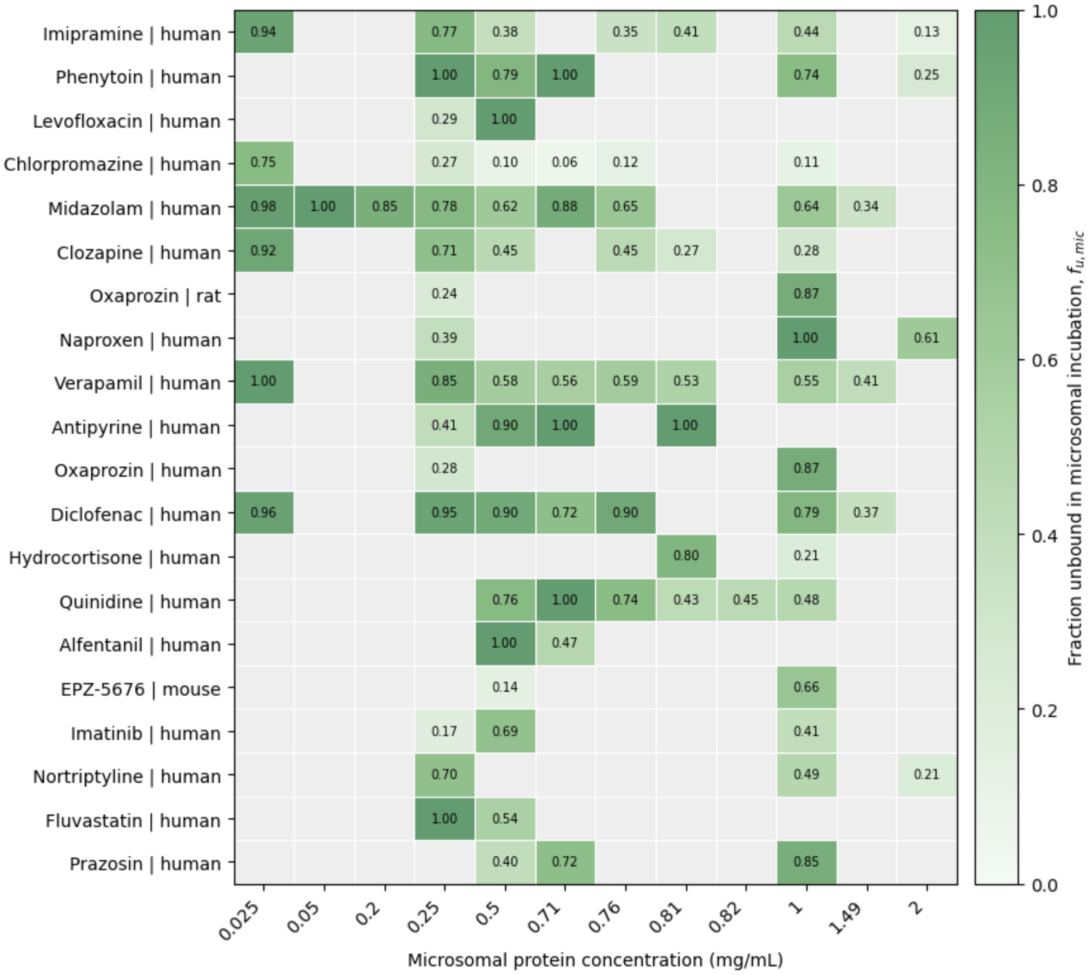


**Figure S1. Variation in microsomal unbound fraction with microsomal protein concentration.** The heatmap shows the 20 compound–species groups with the largest range in fu,mic across at least two protein concentrations in the dataset. Repeated measurements at each concentration were averaged. Cell values and green shading indicate fu,mic; gray cells indicate missing data.

© 2026 MARS, Viva Biotech (Shanghai) Limited. All rights reserved. Visit website at www.vivabiotech.com.

The impact of such assay variability is particularly important for intrinsic-clearance prediction, where experimental uncertainty can approach the magnitude of current model errors. Because CLint is commonly modeled after log10 transformation, modest regression errors may correspond to substantial differences on the original scale. For example, errors of 0.2 and 0.3 log10 units correspond to approximately 1.6- and 2.0-fold differences, respectively [10,11]. Notably, a large-scale multispecies study using more than 200,000 compounds and an ensemble graph neural network reported prospective MAEs of approximately 0.24 to 0.29 log10 units (GMFE, 1.74 to 1.96-fold) across species [12]. Experimental replicate measurements themselves showed considerable variability, with only approximately 77-84% of repeated microsomal CLint measurements falling within twofold. These observations indicate that experimental variability represents an important component of the apparent prediction error and should be considered when interpreting differences between models.

Related effects may also occur in plasma protein-binding assays, where plasma dilution, protein concentration, and nonspecific adsorption to assay materials can influence the measured unbound fraction, particularly for highly bound or lipophilic compounds [13,14]. These experimental variables should therefore be retained where possible and may also be considered as conditioning features during model development. Furthermore, because several assays can report a parameter termed “half-life,” the experimental context must be explicitly distinguished. Plasma stability half-life, microsomal metabolic half-life, and systemic terminal half-life describe fundamentally different processes and should not be treated as interchangeable labels.

For systemic in vivo PK endpoints, a major source of ambiguity arises from distinguishing absolute, apparent, and route-dependent measurements. Absolute oral bioavailability is determined by comparing dose-normalized systemic exposure after oral and intravenous administration. When an intravenous reference is unavailable, absolute bioavailability cannot be directly determined. Likewise, clearance and volume of distribution estimated following oral or other extravascular administration are commonly reported as apparent parameters, CL/F and V/F, because the unknown bioavailability is incorporated into their estimates. These values should not be directly merged with CL or Vd obtained following intravenous administration. Administration route should therefore be explicitly curated and incorporated into endpoint definition or model conditioning before systemic PK data are used for model development. Additionally, terminal half-life requires route-specific interpretation. Following oral administration, slow absorption can govern the terminal concentration–time profile, producing a longer apparent terminal half-life than after intravenous administration. Under such absorption-limited, or flip-flop, kinetics, a prolonged oral terminal half-life does not necessarily indicate slower systemic elimination. Pooling oral and intravenous half-lives can therefore conflate absorption with elimination [15,16]. Administration route, parameter definition, and relevant dosing conditions should therefore be explicitly considered when curating systemic PK endpoints for model development.

© 2026 MARS, Viva Biotech (Shanghai) Limited. All rights reserved. Visit website at www.vivabiotech.com.

#### 2. Additional figures and tables

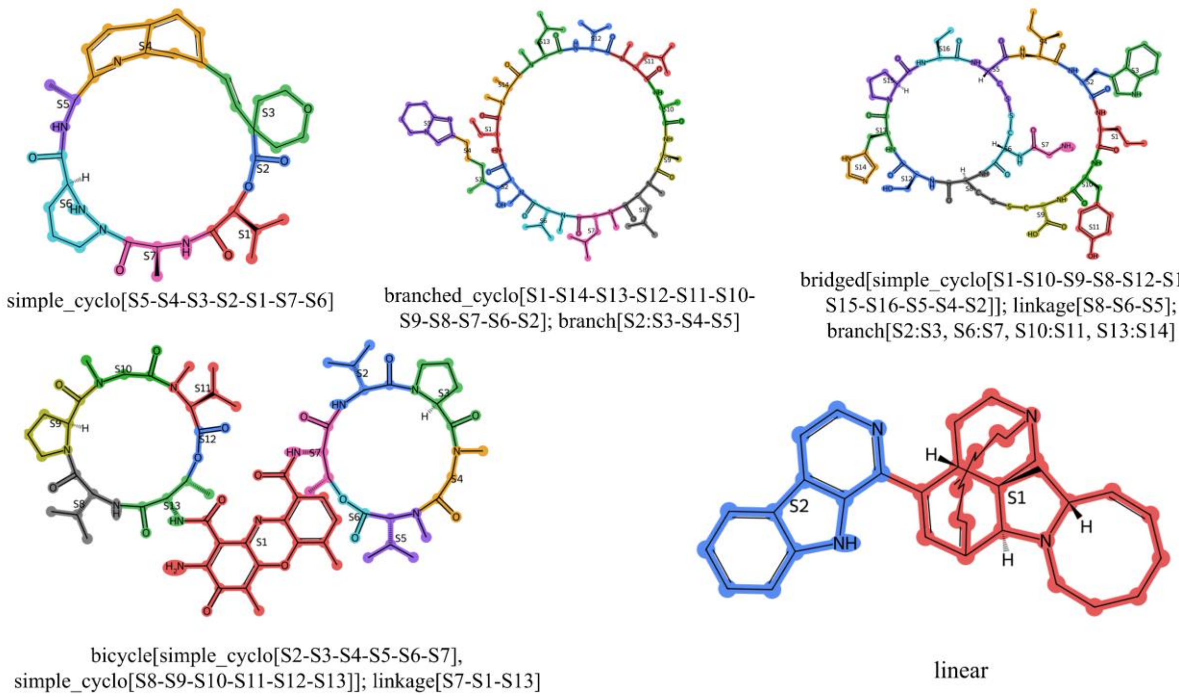


**Figure S2. Representative examples of molecular segmentation and topology annotation.** Examples illustrate simple cyclic, branched cyclic, bridged, bicyclic, and linear representations produced by the segmentation framework. Individual molecular segments are distinguished by color and labeled sequentially (S1, S2, ...), with the corresponding topology annotation shown beneath each structure. The linear example represents the encoding fallback used for structures that cannot be represented by the predefined cyclic topology classes.

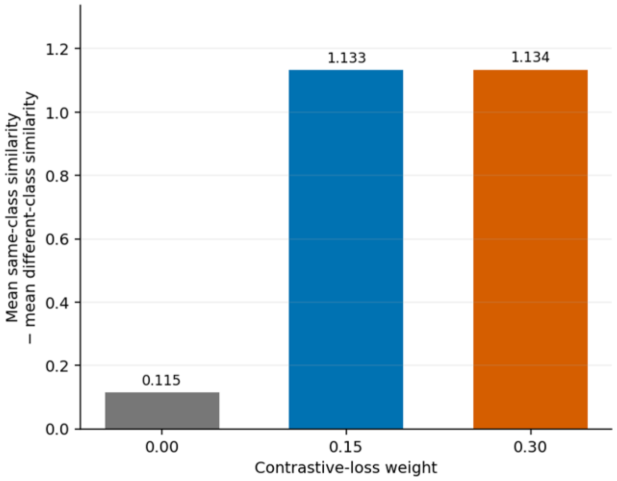


**Figure S3. Effect of contrastive-loss weight on solvent–state representation separation.** Bars show the difference between mean cosine similarity for conformer pairs belonging to the same solvent–state class and that for pairs from different classes. Increasing the contrastive-loss weight from 0 to 0.15 markedly increased class separation, with little additional change at 0.30.

© 2026 MARS, Viva Biotech (Shanghai) Limited. All rights reserved. Visit website at www.vivabiotech.com. 

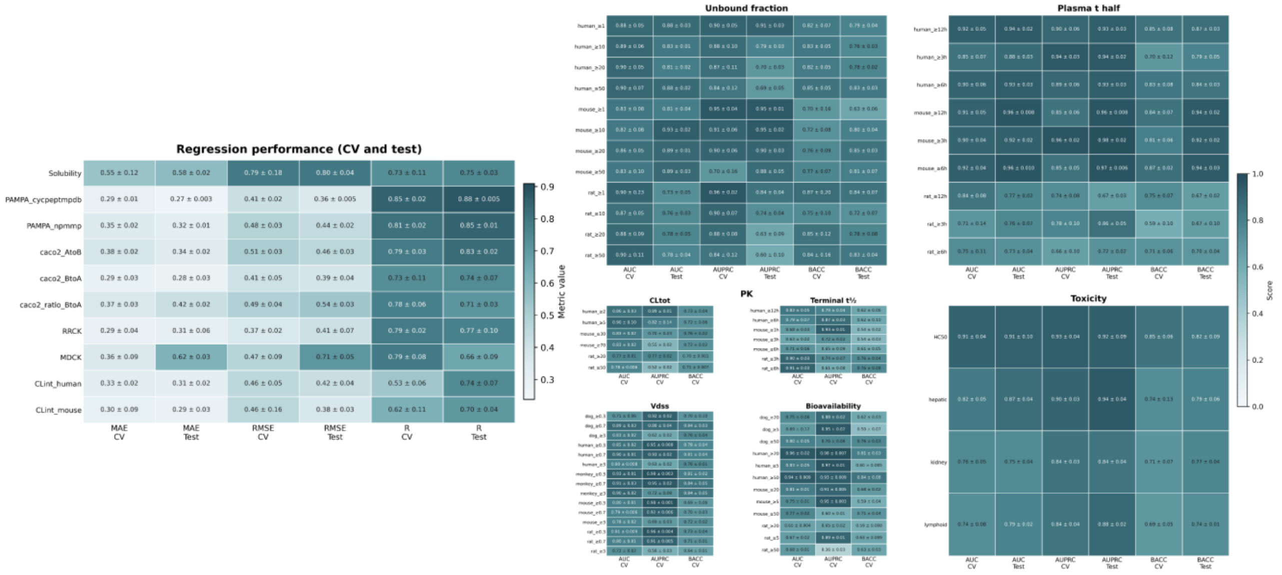


**Figure S4. Cross-validation and held-out test performance of AIO across ADMET endpoints.** Left, regression performance for solubility, permeability, and intrinsic-clearance endpoints, evaluated using MAE, RMSE, and Pearson R. Right, classification performance across plasma unbound fraction, plasma/serum half-life, total clearance (CLtot), terminal half-life, volume of distribution at steady state (VDss), oral bioavailability, and toxicity endpoints, evaluated using AUROC, AUPRC, and BACC. Results are shown for both cross-validation (CV) and held-out test sets; results were described as value ± SD.

© 2026 MARS, Viva Biotech (Shanghai) Limited. All rights reserved. Visit website at www.vivabiotech.com.

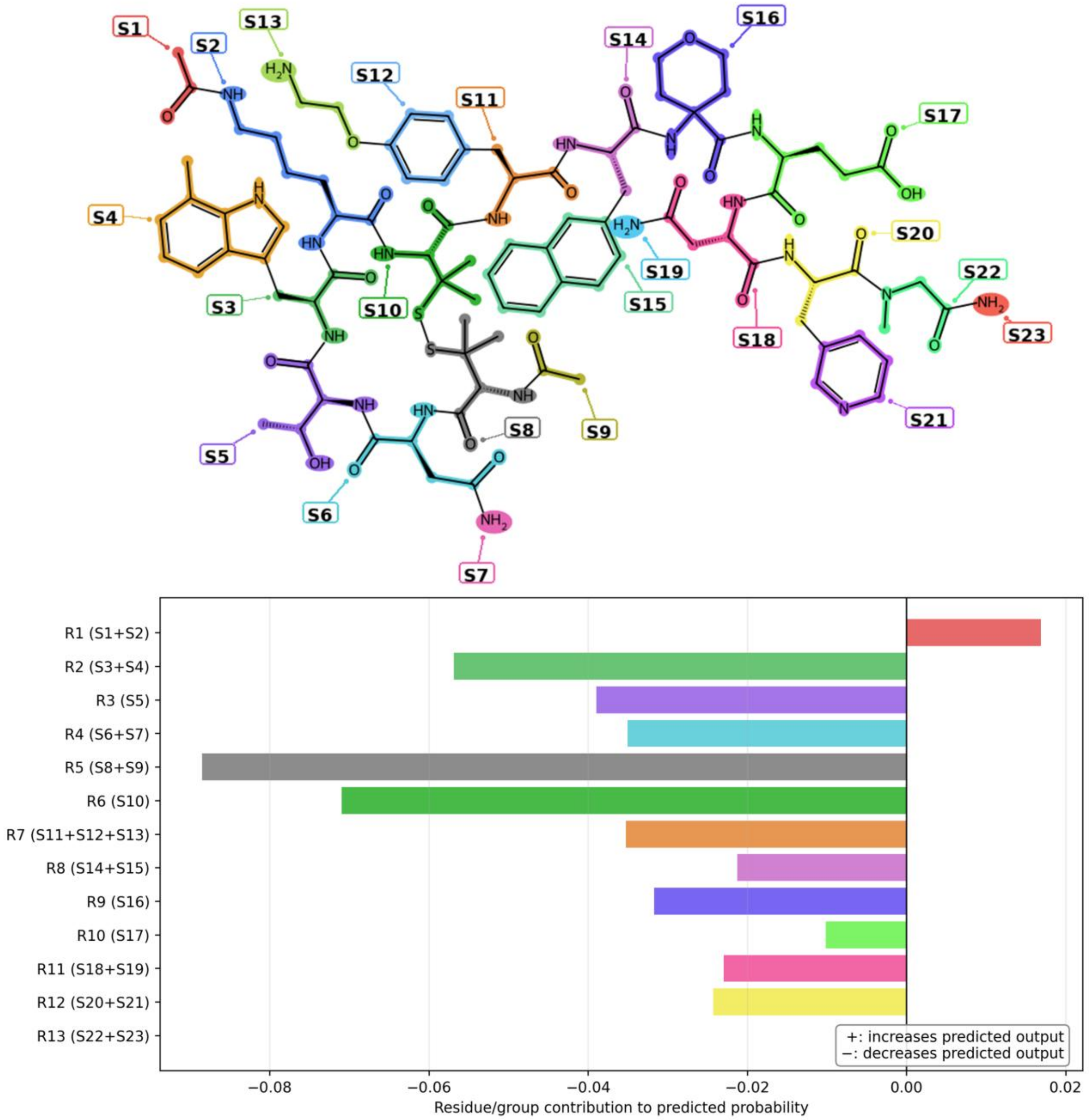


**Figure S5. Detailed segment-level attribution profile for predicted oral bioavailability of icotrokinra.** Top, molecular segmentation of icotrokinra, with individual segments labeled S1–S23. Bottom, contribution of each predefined segment group to the predicted bioavailability probability. Positive values indicate groups that increase the predicted bioavailability output, whereas negative values indicate groups that decrease it. The strongest negative contributions were localized to R5 (S8+S9), R6 (S10), and R2 (S3+S4).

**Table S2. Decision thresholds used to formulate classification tasks across ADMET endpoints.** Thresholds were selected to represent progressively distinct pharmacokinetic regimes and, where applicable, were informed by physiologically relevant reference ranges. Species-specific

© 2026 MARS, Viva Biotech (Shanghai) Limited. All rights reserved. Visit website at www.vivabiotech.com.

thresholds were used for total clearance and intravenous terminal half-life because of interspecies differences in systemic pharmacokinetics.

| Endpoint for classification | Species | Decision thresholds | Rationale |
| --- | --- | --- | --- |
| Unbound fraction | All | 1%, 10%, 20%, 50% | 1%: extreme binding; higher cutoffs: increasing free fraction. |
| Plasma/serum stability | All | 3, 6, 12 h | rapid degradation to sustained stability in plasma/serum |
| Vdss | All | 0.3, 0.7, 3.0 L/kg | 0.3–0.7 L/kg: limited distribution within physiological fluid volumes, >3 L/kg: substantial tissue partitioning. |
| Total clearance | Human | 2, 5 ml/min/kg | Low-clearance range relevant to exposure optimization; 1/3 and 2/3 $Q_H$ for human as physiological references. |
| Total clearance | Rat | 20, 50 ml/min/kg | 1/3 and 2/3 $Q_H$ for rats as physiological references. |
| Terminal half-life (IV) | Human | 6, 12 h | Short to sustained systemic exposure |
| Terminal half-life (IV) | Rat | 3, 6 h | Short to sustained systemic exposure |
| Oral bioavailability (F) | All | 5%, 20%, 50% | <5%: may need further optimization to improve F; > 50%: favorable oral exposure |

**Table S3. Comparison of AIO with conventional regression baselines on the fixed test sets.** Results are reported as MAE/Pearson's (R). kNN and XGB use Morgan fingerprints as molecular inputs, whereas GNN learns representations directly from molecular graphs. Predictions from the cross-validation models were averaged for fixed-test evaluation. PAMPA results are reported separately by data source, and liver microsomal intrinsic clearance (CLint) is reported separately for human and mouse data. The best result for each endpoint and metric is shown in bold.

| Endpoint | AIO | AIO (no pretraining) | GNN | kNN | XGB |
| --- | --- | --- | --- | --- | --- |
| PAMPA_cycpeptmpdb | **0.27 / 0.88** | 0.29 / 0.86 | 0.42 / 0.61 | 0.35 / 0.78 | 0.34 / 0.80 |
| PAMPA_npmmp | **0.32 / 0.85** | 0.32 / 0.85 | 0.45 / 0.75 | 0.41 / 0.76 | 0.35 / 0.85 |
| Solubility | **0.58 / 0.75** | 0.63 / 0.70 | 0.73 / 0.57 | 0.69 / 0.62 | 0.64 / 0.70 |
| Caco-2_AtoB | **0.34 / 0.83** | 0.35 / 0.83 | 0.45 / 0.73 | 0.38 / 0.77 | 0.35 / 0.82 |
| Caco-2_BtoA | **0.28 / 0.74** | 0.29 / 0.74 | 0.37 / 0.49 | 0.34 / 0.58 | 0.34 / 0.50 |

© 2026 MARS, Viva Biotech (Shanghai) Limited. All rights reserved. Visit website at www.vivabiotech.com.

| | | | | | |
|---|---|---|---|---|---|
| Caco-2_ratio_BtoA | 0.42 / 0.71 | **0.39 / 0.77** | 0.57 / 0.42 | 0.48 / 0.60 | 0.45 / 0.67 |
| RRCK | **0.31 / 0.77** | 0.40 / 0.56 | 0.43 / 0.55 | 0.35 / 0.74 | 0.36 / 0.71 |
| MDCK | **0.62 / 0.66** | 0.67 / 0.49 | 0.85 / 0.06 | 0.68 / 0.23 | 0.54 / 0.51 |
| CLint_human | **0.31 / 0.74** | 0.33 / 0.67 | 0.33 / 0.73 | 0.32 / 0.70 | 0.32 / 0.69 |
| CLint_mouse | 0.29 / 0.70 | 0.26 / **0.73** | 0.28 / 0.71 | **0.25** / 0.70 | 0.28 / 0.67 |

Conventional baseline comparisons were restricted to continuous regression endpoints. Classification analyses instead focused on evaluating the effect of pretraining within an identical multi-task architecture across species- and threshold-specific prediction heads (Table S4).

**Table S4. Effect of pretraining on classification performance.** Results are reported as macro-AUROC/macro-BACC, averaged from multiple species and decision thresholds. The better result for each metric is shown in bold.

| Endpoint | AIO (Pretrained) | AIO (No pretraining) |
|---|---|---|
| Unbound fraction | **0.83 / 0.78** | 0.82 / 0.76 |
| Plasma/serum half-life | **0.88 / 0.82** | 0.86 / 0.76 |
| Total clearance | **0.83 / 0.72** | 0.80 / 0.69 |
| Vdss | **0.83 / 0.75** | 0.81 / 0.71 |
| Terminal half-life | 0.78 / 0.64 | **0.80 / 0.66** |
| Oral bioavailability | **0.76 / 0.66** | 0.74 / 0.61 |
| HC50 | **0.91 / 0.82** | 0.84 / 0.74 |
| CC50 | **0.79 / 0.76** | 0.76 / 0.73 |

© 2026 MARS, Viva Biotech (Shanghai) Limited. All rights reserved. Visit website at www.vivabiotech.com.

3. Badée, J. *et al.* Optimization of Experimental Conditions of Automated Glucuronidation Assays in Human Liver Microsomes Using a Cocktail Approach and Ultra-High Performance Liquid Chromatography–Tandem Mass Spectrometry. *Drug Metab. Dispos.* **47**, 124–134 (2019).
4. Katneni, K. *et al.* Using Human Plasma as an Assay Medium in Caco-2 Studies Improves Mass Balance for Lipophilic Compounds. *Pharm. Res.* **35**, 210 (2018).
5. Wang, N.-N. *et al.* ADME Properties Evaluation in Drug Discovery: Prediction of Caco-2 Cell Permeability Using a Combination of NSGA-II and Boosting. *J. Chem. Inf. Model.* **56**, 763–773 (2016).
6. Winiwarter, S. *et al.* Prediction of Fraction Unbound in Microsomal and Hepatocyte Incubations: A Comparison of Methods across Industry Datasets. *Mol. Pharm.* **16**, 4077–4085 (2019).
7. Chen, S., Prieto Garcia, L., Bergström, F., Nordell, P. & Grime, K. Intrinsic Clearance Assay Incubational Binding: A Method Comparison. *Drug Metab. Dispos.* **45**, 342–345 (2017).
8. Krumpholz, L., Klimczyk, A., Bieniek, W., Polak, S. & Wiśniowska, B. Data set of fraction unbound values in the in vitro incubations for metabolic studies for better prediction of human clearance. *Database* **2024**, baae063 (2024).
9. Koopmans, A. B., Braakman, M. H., Vinkers, D. J., Hoek, H. W. & Harten, P. N. van. Meta-analysis of probability estimates of worldwide variation of CYP2D6 and CYP2C19. *Transl. Psychiatry* **11**, 141 (2021).
10. Iwata, H. *et al.* Predicting Total Drug Clearance and Volumes of Distribution Using the Machine Learning-Mediated Multimodal Method through the Imputation of Various Nonclinical Data. *J. Chem. Inf. Model.* **62**, 4057–4065 (2022).
11. Keefer, C. E. *et al.* The Comparison of Machine Learning and Mechanistic In Vitro–In Vivo Extrapolation Models for the Prediction of Human Intrinsic Clearance. *Mol. Pharm.* **20**, 5616–5630

© 2026 MARS, Viva Biotech (Shanghai) Limited. All rights reserved. Visit website at www.vivabiotech.com.

(2023).

© 2026 MARS, Viva Biotech (Shanghai) Limited. All rights reserved. Visit website at www.vivabiotech.com.
